# The Clinical Imperative for Inclusivity: Race, Ethnicity, and Ancestry (REA) in Genomics

**DOI:** 10.1101/317800

**Authors:** A.B. Popejoy, D.I. Ritter, K. Crooks, E. Currey, S.M. Fullerton, L.A. Hindorff, B. Koenig, E.M. Ramos, E.P. Sorokin, H. Wand, M.W. Wright, J. Zou, C.R. Gignoux, V.L. Bonham, S.E. Plon, C.D. Bustamante, The Clinical Genome Resource (ClinGen) Ancestry and Diversity Working Group (ADWG)

**Affiliations:** Stanford University; Baylor College of Medicine; University of Colorado, Anschutz Medical Campus; National Human Genome Research Institute (NHGRI); University of Washington; University of California at San Francisco (UCSF)

**Keywords:** Ancestry, Race, Ethnicity, Populations, Genomics, Diversity

## Abstract

The Clinical Genome Resource (ClinGen) Ancestry and Diversity Working Group highlights the need to develop guidance on race, ethnicity, and ancestry (REA) data collection and use in clinical genomics. We present quantitative and qualitative evidence to characterize: 1) acquisition of REA data via clinical laboratory requisition forms, and 2) information disparity across populations in the Genome Aggregation Database (gnomAD) at clinically relevant sites as determined by variants in ClinVar. Our requisition form analysis showed substantial heterogeneity in clinical laboratory ascertainment of REA, as well as marked incongruity among terms used to define REA categories. There was also striking disparity across REA populations in the amount of information available about variants at clinically relevant sites in gnomAD. European ancestral populations constituted the majority of observations (55.8%), allele counts (59.7%), and private alleles (56.1%) in gnomAD at 550 loci with “pathogenic” and “likely pathogenic” expert-reviewed variants in ClinVar. Our findings highlight the importance of implementing and supporting programs to increase diversity in genome sequencing and clinical genomics, as well as measuring uncertainty around population-level datasets that are used in variant interpretation. Finally, we suggest the need for a standardized REA data collection framework to be developed and adopted across clinical genomics.

## Background

Global efforts such as the National Institutes of Health (NIH)-funded Clinical Genome Resource (ClinGen)^1^ seek to greatly increase curation activities and develop resources for clinicians and scientists to study the relationship among diseases, genes and variants. As such, it is critical to understand how race, ethnicity, and genetic ancestry (REA) should be ascertained, communicated, and applied in clinical genomics. Clinical sequencing is now being routinely used for diagnosis, prognosis, and treatment. This has spilled over into the direct-to-consumer (DTC) market, where ancestry and disease products are expanding in popularity among the general public in the United States (Ramos & Weissman, 2018; Roberts et al, 2017). As a consequence of tens of millions of people already having access to their genetic data, physicians are faced with diverse populations of patients asking about the clinical significance of their genetic information, and this trend will continue to grow in the future. The collection of REA data from patients is often implemented in self-reported contexts (Bonham et al., 2017), for which there are no established best practices, and challenges include complexities of ancestral admixture and international differences in the use of racial, ethnic, and ancestral categories (Morales et al., 2018; Smart et al., 2017; Yu et al., 2012; Travassos & Williams, 2004).

At present, the “precision” of precision medicine is based on a foundation of evidence mostly from European ancestry populations (Landry et al., 2018; Morales et al., 2018, Popejoy & Fullerton, 2016). Patients of non-European ancestry are more likely to receive ambiguous genetic test results (variants of unknown or uncertain significance, VUS) (Caswell-Jin et al., 2018), false positive diagnoses (Manrai et al., 2016), and false negative diagnoses in the absence of a robust genomic evidence-base (Sorokin et al., *In Press*) and variant reclassification over time (Hiatt et al., 2018). Thus, investigators and professionals in the clinical and scientific research communities have a responsibility to increase diversity in clinical and research sequencing as well as to understand and harmonize the presentation and utilization of REA in clinical genomics. In this paper, we address effects on clinical genomics from the lack of diversity in databases and a lack of clarity in terms used to describe people and populations.

The ClinGen Ancestry and Diversity Working Group (ADWG) formed in November 2017 to improve our understanding of the role of genomic diversity across populations in a clinical genomics setting. Our work includes contributing to the knowledge-base around scientifically and culturally complex issues of self-reported race, ethnicity, geographic and genomic ancestry, particularly in the context of interpreting variants identified via clinical sequencing. The ADWG is multi-disciplinary, comprising investigators and clinicians from genomic medicine, bioethics, health disparities, public health, genetic epidemiology, and statistical and population genetics. The diverse expertise of our members, our collaboration with other ClinGen working groups, and the strong commitment of the National Human Genome Research Institute (NHGRI) to promoting diversity and research on ancestry (Hindorff et al., 2018; Manolio et al., 2017; Green et al., 2011) results in the ADWG being well-poised to tackle such complex issues.

We describe here the initial phase of ADWG efforts, focusing on pilot studies that evaluate two core areas in clinical genomics: 1) Ascertainment of patient race, ethnicity, and ancestry; and 2) Information disparity across populations for genetic variants that are clinically relevant. We are also surveying clinical genomics practitioners to investigate knowledge, beliefs, and use of REA in variant curation and clinical applications. Our findings are meant to provide a baseline understanding in the clinical genomics community about how race, ethnicity, and ancestry are conceptualized and utilized in current practice, with the future aim of developing standards and recommendations to guide how these concepts inform sequence variant interpretation and subsequent decision making by clinical practitioners.

The striking information disparity between European ancestry populations and all others in terms of available data underscores the importance of continued efforts to diversify the genomic evidence-base. It also strongly suggests that uncertainty around allele frequency estimates is likely to be much higher for non-European ancestry populations, and downstream analyses such as disease risk predictions may be less accurate for individuals from these populations as well.

## Methods

### Clinical Ascertainment

In order to establish a baseline understanding of how race, ethnicity, and ancestry (REA) are ascertained in a clinical genomics setting, we analyzed genetic test requisition forms from each of ten clinical laboratories that have contributed the most variant interpretations to ClinVar and have analyzed >2 genes and conditions. For this analysis, *Clinvar Miner*^2^ was accessed on 3/2/18 to identify a list of the top ten submitters to ClinVar filtering on criteria for collection method (clinical testing) and submission review status (evidence provided with the assertion from at least one submitter). Laboratory websites were manually accessed to download requisition forms (RFs) between 3/2/18 - 3/6/18. Preference was given to prenatal or carrier testing, hereditary cancers, and general RFs. In one case, laboratory RFs were not available online, but were located through websites of other laboratories that use them. In another case, the laboratory RFs were not available from the laboratory website to non-clinicians, so the lab sent us a patient history form, which asks the same REA question as their test RFs. However, most laboratory RFs were available online and provided consistent categories across forms; in total, we analyzed forms from ten different labs who were among the top submitters to ClinVar and had RFs publicly available.

If consistency in the RF options for REA was observed among at least three RFs, then one was chosen as the “representative” RF for that laboratory. Only one of the ten laboratories analyzed provided different REA options to choose from across the RFs viewed, and in that case the RF with the maximal number of REA options was used. Screenshots of the relevant questions and possible answers were captured and logged in a working document, examples of which can be viewed in the Appendix. REA options provided on different forms were recorded in an Excel file, in addition to labels used to designate that section on the RF (such as “Ancestry” or “Race and Ethnicity”), which differed across forms. Finally, we tabulated the number of times each specific REA option was observed in any RF and associated it with a representative category from the population ontology used by MedGen^3^.

### Information Disparity

To create the variant set for an information disparity analysis, we obtained a .csv file of all unique GRCh37 (hg19) “pathogenic” or “likely pathogenic” entries in ClinVar for single-nucleotide variants (SNVs) that have been reviewed by an expert panel (N=1661). We further restricted the analysis to genes with at least 10 variants observed in ClinVar; 96% of the initial variants at unique sites were found within these eight genes (N=1585 variants). Population-level data including allele counts, number of private alleles, and numbers of observations were obtained from population groupings in gnomAD (accessed April 2018): African/African-American (AFR), Ashkenazi Jewish (ASJ), East Asian (EAS), non-Finnish European (NFE), Finnish (FIN), Latino or admixed American (AMR), and South Asian (SAS). We further restricted our analysis to variants identified from ClinVar that had corresponding entries in gnomAD exomes or genomes (N=550) to determine the level of information disparity at those sites.

## Results

### Ascertainment Heterogeneity and Ambiguity

Our analysis of requisition forms (RFs) revealed considerable heterogeneity across clinical laboratories in the way REA data are ascertained. Out of questions asked on RFs from ten different laboratories, only one asked an open-ended question with a blank “Geoancestry / Ethnicity” field. All others provided pre-determined selections in a multiple-choice or best-choice response format. Across forms from the same clinical lab, the questions and possible responses were usually identical, regardless of the type of test. Among labs, however, the combination and number of pre-set selections varied widely. Some specific race and ethnicity categories were consistent across most forms, such as “African American” (9/10), “Ashkenazi Jewish” (8/10), “Asian” (8/10) and “Hispanic” (9/10). Variable REA terms were used to describe European-ancestry populations, such as “White” (2/10) and “Caucasian” (7/10). Many forms also offered more granular European categories based on geographic origin: “Northern European, e.g. British, German”, “Southern European e.g. Italian, Greek”, “Western European”, and “Eastern European”, for example. Similarly, categories based on geography were used to define various Asian populations on one form: “East Asian e.g. Chinese, Japanese”, “South Asian e.g. Indian Pakistani”, and “Southeast Asian e.g. Filipino, Vietnamese”. In contrast, large continents and highly diverse populations were described using racial categories such as “Black/African American” (1/10), “African or African American” (1/10), “Native American” (6/10), and “Indigenous” (1/10) with no geographic specificity. Clinical laboratory RFs thus varied in how they identified populations, which included continental ancestral population, country of origin, self-identified race, and ethnicity, often without distinguishing between the different types of population identifiers.

In order to organize these heterogeneous REA labels into broad categories, we identified the MedGen population codes used in ClinVar and mapped each of the terms to one of those categories (Table 1). Among the nine RFs that offered pre-set REA selections, there were 38 unique terms used to describe roughly eight large continental populations or ancestral designations found in MedGen, and the majority of those terms (28/38) were used only once on a single RF. Ten of the unique descriptors observed on RFs are variations of a European ancestry population, typically a specific geographic region. In contrast, there were only three very similar descriptors available for African Americans, none of which refers to any geographic region, and all three of which included the term “African American”. There were also three instances of terms used to describe Ashkenazi Jewish ancestry or ethnicity, and all three of them included “Jewish”. Finally, there were several terms used to designate groups that did not clearly correspond to any MedGen categories, including “Caribbean”, “Central/South American”, “French Canadian or Cajun”, and “Jewish-Sephardic”, each of which were each seen only once, and “Middle Eastern”, which was seen four times.

**Table 1.** Terms used to describe human racial, ethnic, and ancestry groups on requisition forms for highly productive ClinVar-submitting laboratories. Each term has a number in parentheses after it, which represents the number of forms on which the corresponding term was observed. Theme headers are the MedGen categories used as population descriptors in ClinVar, and terms are matched to these categories.

|  |  |
| --- | --- |
| **African American; MedGen:C0085756** | **Hispanic Americans; MedGen:C0019576** |
| African American (6) | Hispanic (9) |
| African or African American (1) | **Native Hawaiian or Other Pacific Islander; MedGen:C1513907** |
| Black/African American (2) | Native Hawaiian or other Pacific Islander (1) |
| **American Indian or Alaska Native; MedGen:C1515945** | Pacific Islander (3) |
| American Indian / Native American (1) | **Chinese; MedGen:C0152035 / South East Asian; MedGen:C0238697** |
| Native American (5) | Asian (7) |
| Indigenous (1) | Asian/Pacific Islander (1) |
| **Ashkenazi Jew; MedGen:C0337704** | E. Indian (1) |
| Ashkenazi Jewish (7) | East Asian e.g. Chinese, Japanese (1) |
| Jewish (1) | South Asian e.g. Indian Pakistani (1) |
| Jewish-Ashkenazi (1) | Southeast Asian e.g. Filipino, Vietnamese (1) |
| **Caucasians; MedGen:C0043157 / European Caucasoid; MedGen:C0682087** | **Mixed Ethnic Group; MedGen:C0682086** |
| Caucasian (5) | Other/Mixed Caucasian (1) |
| Caucasian/NW European (1) | **None / Unspecified** |
| Eastern European (1) | Adopted (1) |
| Mediterranean (1) | Other: _____ (8) |
| Northern European (1) | Unknown (1) |
| Northern European e.g. British, German (1) | **[Other: No MedGen Category Clearly Applies]** |
| Portuguese (1) | Caribbean (1) |
| Southern European e.g. Italian, Greek (1) | Central/South American (1) |
| Western European (1) | French Canadian or Cajun (1) |
| White (1) | Jewish-Sephardic (1) |
| White/Caucasian (1) | Middle Eastern (4) |

Overall, no two clinical laboratories provided the same descriptive categories to designate a group or population on their RFs. Nor was there consistency in the way laboratories described the category of information being sought about REA. Metadata headers (for the REA section of lab RFs) included terms such as “Ancestry”, “Ethnicity”, “Race and Ethnicity”, or blank (pre-selected race and ethnicity options, field was not named). Table 1 shows all of the terms used to describe various groups and populations, organized by their corresponding MedGen category as determined by the authors. We acknowledge that there may be differences of opinion with how each of these terms is associated (or not) with the MedGen categories, and this presentation is not meant to suggest this is the best framework for mapping those associations. Rather, it is intended to organize the diverse categories we observed and clearly show the heterogeneity and ambiguity with which people are grouped in a clinical genomics setting, illuminating the importance of efforts to improve the ascertainment of self-reported race and ethnicity.

### European-Biased Clinical Genomic Evidence

It is well established that genomic databases on which clinical variant interpretations are made are biased toward European ancestry, but the amount of information from different REA populations specific to clinically relevant sites is less clear. We conducted an analysis to quantify the information disparity across populations in two databases that are typically used for clinical variant curation: ClinVar (Landrum et al., 2014) and the Genome Aggregation Database (gnomAD) (Lek et al., 2016).

Of the 1805 variants that have been reported to ClinVar and determined by an expert panel to be “pathogenic” or “likely pathogenic”, 1661 of these are found at unique sites in the genome. Of these unique sites, 1585 (96%) are found in eight genes, as a limited number of expert panels have completed the ClinVar/ClinGen approval process [*BRCA1* (388), *BRCA2* (473), *CFTR* (154), *MLH1* (232), *MSH2* (195), *MSH6* (72), *MYH7* (44), and *PMS2* (27)]. Among the 1585 clinically relevant sites in these genes, 550 (35%) had a corresponding variant entry in gnomAD, not surprisingly, as most pathogenic variants represent rare alleles in population databases (Lek et al. 2016). We determined what proportion of total observations, allele counts, and private (population-specific) alleles in a particular gene were attributed to each population and illustrate the resulting distribution in Figure 1. Our primary finding is that all three of these (non-independent) metrics show the majority of information is from European-ancestry individuals.

**Figure 1.**
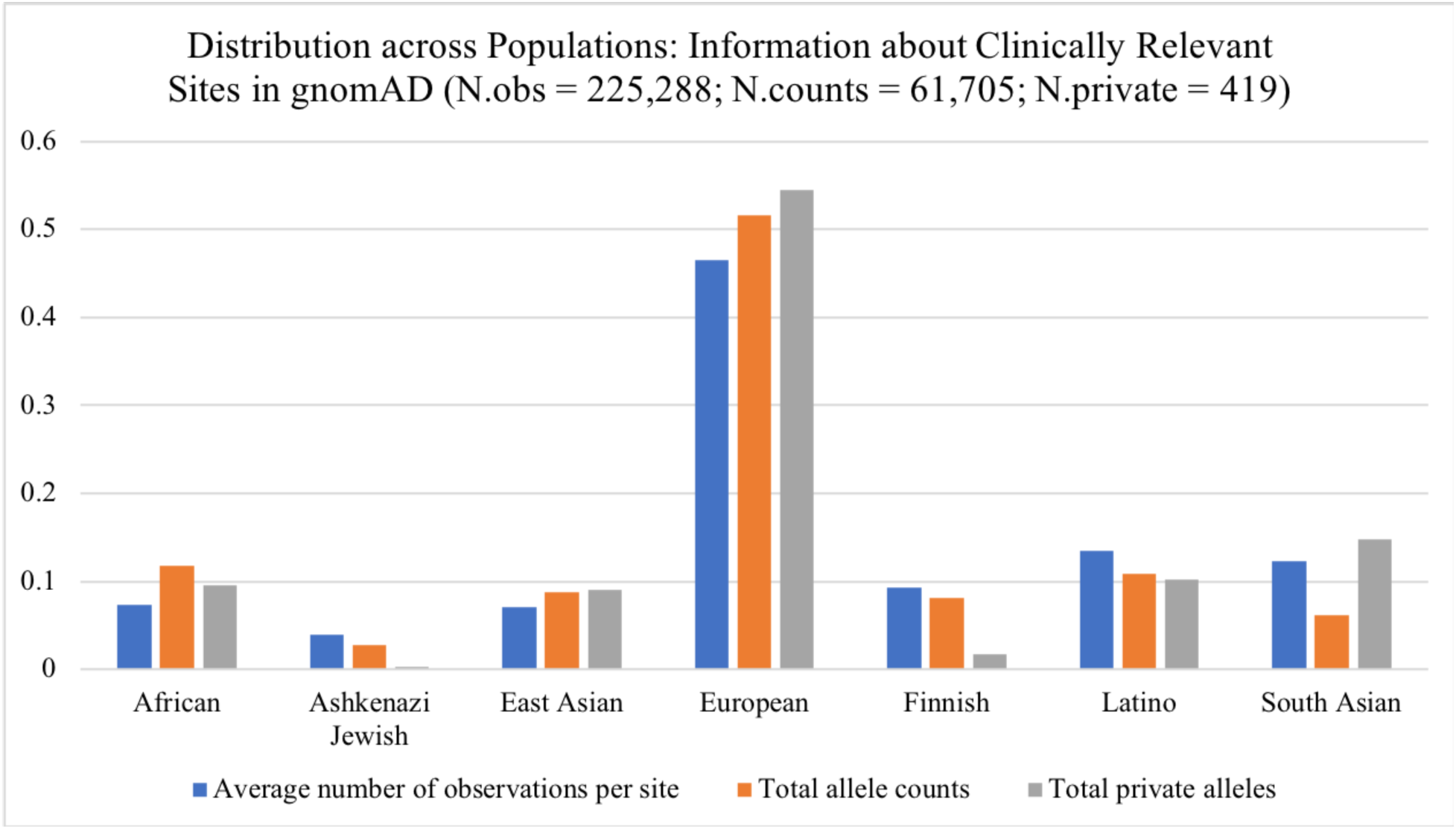
Information Disparity at Clinically Relevant Sites across Populations. Bar graph shows the distribution of information that is available for different ancestral populations in gnomAD, for variants in ClinVar that are annotated by expert review as “pathogenic” or “likely pathogenic”. The percentage of average total observations (blue, N=225,288) is 7.4% African (AFR), 4% Ashkenazi Jewish (ASJ), 7.1% East Asian (EAS), 46.5% European (NFE), 9.3% Finnish (FIN), 13.5% Latino (AMR), 12.3% South Asian (SAS). The percentage of total allele counts (orange, N=61,705) observed across populations is 11.8% AFR, 2.7% ASJ, 8.1% EAS, 51.6% NFE, 8.2% FIN, 10.9% AMR, and 6.2% SAS. Private alleles are those found only in a single population (gray, N=419); the percentage of all private alleles observed was 9.5% in AFR, 0.2% ASJ, 9.1% EAS, 54.4% NFE, 1.7% FIN, 10.3% AMR, and 14.8% SAS.

Overall, roughly 50% of all information about clinically relevant variants (number of observations at each site, observed allele counts, and alleles that are observed in only one population, ‘private alleles’) came from non-Finnish European (NFE) ancestry individuals. It is worth noting that closer to 60% of all information can be attributed to “white” racial groups, including NFE, FIN, and ASJ. Every population except for NFE contributed an additional ~10% of the total information. Latino or admixed Americans (AMR) had the highest number of total observations among non-Europeans (13.5%) while EAS had the lowest (7.1%). After NFE, SAS had the highest proportion of population specific alleles (14.8% that are specific to South Asia) while ASJ and FIN had noticeably fewer (0.2% Ashkenazi Jewish and 1.7% Finnish, respectively). All comparisons are highly significant due to the depth of sequencing available in gnomAD (pairwise comparisons of counts:total observations, all p<1e-4).

## Discussion

Our findings from both the clinical laboratory requisition form analysis and the quantification of information disparity across populations at clinically relevant genetic loci reveal an underlying deficiency in clinical genomics research and implementation in diverse populations. How clinical labs ascertain information about REA is so highly variable that large National Institutes of Health (NIH)-supported consortia such as the Clinical Sequencing Evidence-Generating Research (CSER)^4^ are actively engaged in discussions to harmonize these measures across clinical sites. Indeed, the marked heterogeneity we found here in REA ascertainment by clinical laboratories may have far-reaching downstream effects for research and clinical practice in the era of genomic technology. There is a growing literature describing the downstream effects of a European-biased evidence-base for genomic medicine (Landry & Rehm, 2018). For example, a variant that had been determined pathogenic based on observations in mostly European ancestry individuals led to pervasive false positive diagnoses of African Americans for hypertrophic cardiomyopathy (Manrai et al., 2016). Subsequent invasive treatments were determined to have been unnecessary after the variant was observed in a sufficient number of healthy African Americans, then re-designated as benign. As demographics shift toward a more diverse population, clinical genomics must focus on ways to improve variant interpretation for individuals of admixed and non-European ancestry.

Our results showed prevalent information disparity between populations at clinically relevant sites, and only one-third of sites in the genome with “pathogenic” and “likely pathogenic” variants in ClinVar have readily available population-level data. This has important implications for variant curation and interpretation using the ACMG Guidelines (Richards et al., 2015) as several of their criteria rely on allele frequencies. One criterion for moderate evidence of pathogenicity is specifically related to the absence of a variant from “population databases” (PM2 evidence code). However, absence of a variant is difficult to distinguish from incomplete sampling across all underrepresented populations in genomic databases (currently applies to all non-European ancestry groups). Also, pathogenic variants tend to be rare, so many more observations from diverse groups in large population databases will be necessary. Further complicating the issue of allele frequency data for use in clinical genomics is the nomenclature incongruity between requisition form REA collection and major control database populations (i.e. “Asian” on clinical laboratory RFs vs. “South Asian” or “East Asian” in ExAC and gnomAD). Such inconsistencies across population labels make it challenging to directly relate variant information in the clinic to population allele frequencies. Equitable sampling across all geographic regions and REA groups would begin to alleviate these complications, particularly as they relate to the PM2 evidence code for variant interpretation, since absence of a variant from “population databases” would be more relevant.

The issue of whether and how to categorize ancestral populations in clinical genomics for improved healthcare, in laboratory requisition forms and online public sequencing repositories, is an admittedly controversial topic. Some argue that whole-genome and whole-exome sequencing will alleviate the need to ascertain self-reported REA measures, since genetic ancestry can be directly inferred from DNA; however, disease outcomes vary between racial and ethnic groups to a larger extent than can be explained by genetic differences and this needs to be accounted for in genomic medicine (Burchard et al., 2003). Thus, carefully collecting these data is necessary, and conceptualizing the various meanings of “race”, “ethnicity”, and “ancestry” is an active and ongoing debate (Hunt & Megyesi 2008). Historical and social science research documents how REA terms have varied meaning and are fluid constructs (Yudell, 2014; Smedley & Smedley, 2012; Race, Ethnicity, and Genetics Working Group, 2005). Terms such as “Hispanic,” which was present on all requisition forms that offered an REA choice, are overall not particularly descriptive, despite its usage as an ethnicity metric in the U.S. Census. Additionally, terms such as “Adopted”, “Other/mixed Caucasian”, “Unknown”, or “Indigenous” may not be at all informative for clinical variant interpretation. Categories describing smaller geographic regions, religious groups, or other genetically isolated populations (“French Canadian/Cajun”, “Ashkenazi Jewish”, etc.) may help raise awareness of population bottlenecks and can inform variant interpretation of unique allele frequencies due to founder effects or genetic drift. Conversely, whether large geographic/continental regions such as “African”, “Asian”, “Western European”, and “South/Central American” have clinical utility as population designators is largely unexamined in the clinical genomics space.

Social and political issues are prevalent in population identity, especially for groups that have been traditionally marginalized and/or subjected to discrimination. For example, “Native American” may be used colloquially but on official documents, “American Indian or Alaska Native” is typically the preferred terminology, since this designation is used in all treaties with the U.S. government. Likewise, in anecdotal accounts, patients have reported hesitation to self-identify as “Jewish”, “Ashkenazi” or “Sephardic” because such labels have been used for political persecution. How this information should be used in clinical genomics needs further research. Deciding the most optimal format and terminology for research purposes and clinical applications as well as harmonizing across laboratories, online repositories of genetic data and sequencing projects will require a dedicated effort across public and private sectors.

Our analysis of information disparity at pathogenic sites substantiated the necessity of increasing diversity in genome sequencing. Historically, methodologies and resources in genomics research have been designed for and implemented on mostly white, European-ancestry individuals, and this legacy is perpetuated in current research. Some methodologists are working to leverage unique characteristics of ancestrally diverse genomes to promote discovery (Park et al., 2018; Wojcik et al., 2017; Aschard et al., 2015), and others are exploring effective incentives to sequence more diverse populations. Recently, national efforts are underway to increase genomic diversity in clinical sequencing and research, including NIH-supported All of Us^5^, CSER^6^, Genome Sequencing Program (GSP)^7^, Population Architecture Using Genomics and Epidemiology (PAGE) (Wojcik et al., 2017; Matise et al., 2011) [National Human Genome Research Institute, NHGRI], and Trans-Omics for Precision Medicine (TOPMed)^8^ [National Heart, Lung, and Blood Institute, NHLBI]. Additionally, attempts to standardize ancestry categories for research have been undertaken, such as *Ancestro* for the Catalog of Genome-Wide Association Studies (GWAS Catalog) (Morales et al., 2018). If adopted, existing frameworks will need to be harmonized and built upon in the development of standard REA categories for genomic databases, and other applications in clinical genomics such as electronic health records (EHRs) and laboratory requisition forms.

Additionally, there are deeply entrenched systems that inhibit broader inclusion of diverse populations, such as structural and institutional racism (Williams et al., 2013), historical misuse of data (Garrison, 2013) and subsequent mistrust of the biomedical research community (Ferrera et al., 2015; Gamble, 1997). As such, there is an inherent tension between the need to acknowledge past harms to communities of color and encouraging participation in genomics research, as well as data sharing, to advance the ability of precision medicine to be equitably distributed and effective in all REA populations. Addressing racial and ethnic disparities in genome interpretability requires sharing of data across millions of sequenced genomes and exomes with rich metadata including context of sequencing (for care, research, DTC, etc.) and REA labels. NHGRI-funded research sequencing efforts are a start, since they promise hundreds of thousands of ethnically diverse exomes and genomes and resulting population allele frequency resources by 2021. Meanwhile, millions of genomes, exomes, and large panel tests are likely to be sequenced in the context of clinical care. Therefore, continued development of data standards and secure sharing protocols for clinical genomics are critical for ameliorating population-level disparities in clinical genomics. Additionally, investments in public health interventions and community resources must be prioritized in parallel to genomic sequencing and database development, to address disparities in disease risk and health outcomes that are due to environmental inequities in underserved populations.

As scientists and clinicians, we need to carefully balance the utility of REA information to improve research and clinical care while not reifying harmful, unfounded beliefs about biological differences between racial groups (Fitzgerald, 2014; Roberts, 2011). Race, ethnicity, and ancestry impact health risks and outcomes, as in certain diseases like asthma, kidney, heart disease, and diabetes (Torgerson et al., 2011; Parsa et al., 2013; Manrai et al., 2016, Landry & Rehm, 2018; SIGMA Consortium & Estrada et al., 2014) as well as pharmacogenomics (Ramamoorthy et al., 2015). However, determining the degree to which genetic and/or shared environmental factors (each of which are differently associated with REA measures) influence complex traits is difficult, and will take both genomic and public health solutions to elucidate. The challenge for the clinical genomics community is to determine how to responsibly usher in the era of precision medicine for all patients (Bonham et al., 2016).

## Conclusion

Inclusion of diverse patients and research participants in clinical genomics is a critical piece of addressing the information disparity in our clinical genomic evidence-base; however, development and standardization of genetic ancestry as well as self-reported race and ethnicity measures are also key components. While large-scale whole-genome and -exome sequencing may someday fully elucidate the genomic architecture of diverse ancestral populations, genomic data alone do not account for population-level differences in disease outcomes, even for classic Mendelian traits. Furthermore, these technologies do not address multi-factorial (social, economic, environmental, etc.) components of complex disease etiology. This is where carefully determined REA measures can provide additional layers of understanding of disease-gene and -variant associations, and they must be harmonized across large-scale research consortia, public and private clinical laboratories, and clinical points of care. Future research should provide guidance on the implications of uncertainty around allele frequency estimates in diverse populations; and in the meantime, those involved in variant curation and interpretation efforts using population-level data should be mindful of information disparities across populations. Ultimately, the clinical genomics community must agree on how to appropriately handle the collection and use of race, ethnicity and ancestry, as well as the utility of this information. The ClinGen ADWG is called upon by NHGRI to provide guidance on these issues, and we welcome others to co-create solutions with us.

## Acknowledgments

Additional members of the ClinGen Ancestry and Diversity Working Group (ADWG) who are not listed as authors include: Berg, J., Dwight, S., Kenny, E., Lee, S.S., Ormond, K.E., Oh, S., Prahbu, S., Rivas, M., Shields, A.E., Thornton, T., and Williams, D., many of whom participated in conversations that led to the development of these projects and/or provided minor edits on the manuscript. This work was supported by a National Human Genome Research Institute (NHGRI) grant funding the Clinical Genome Resource (ClinGen) jointly awarded to Stanford University and Baylor College of Medicine.

## Appendix

**Figure A1.**
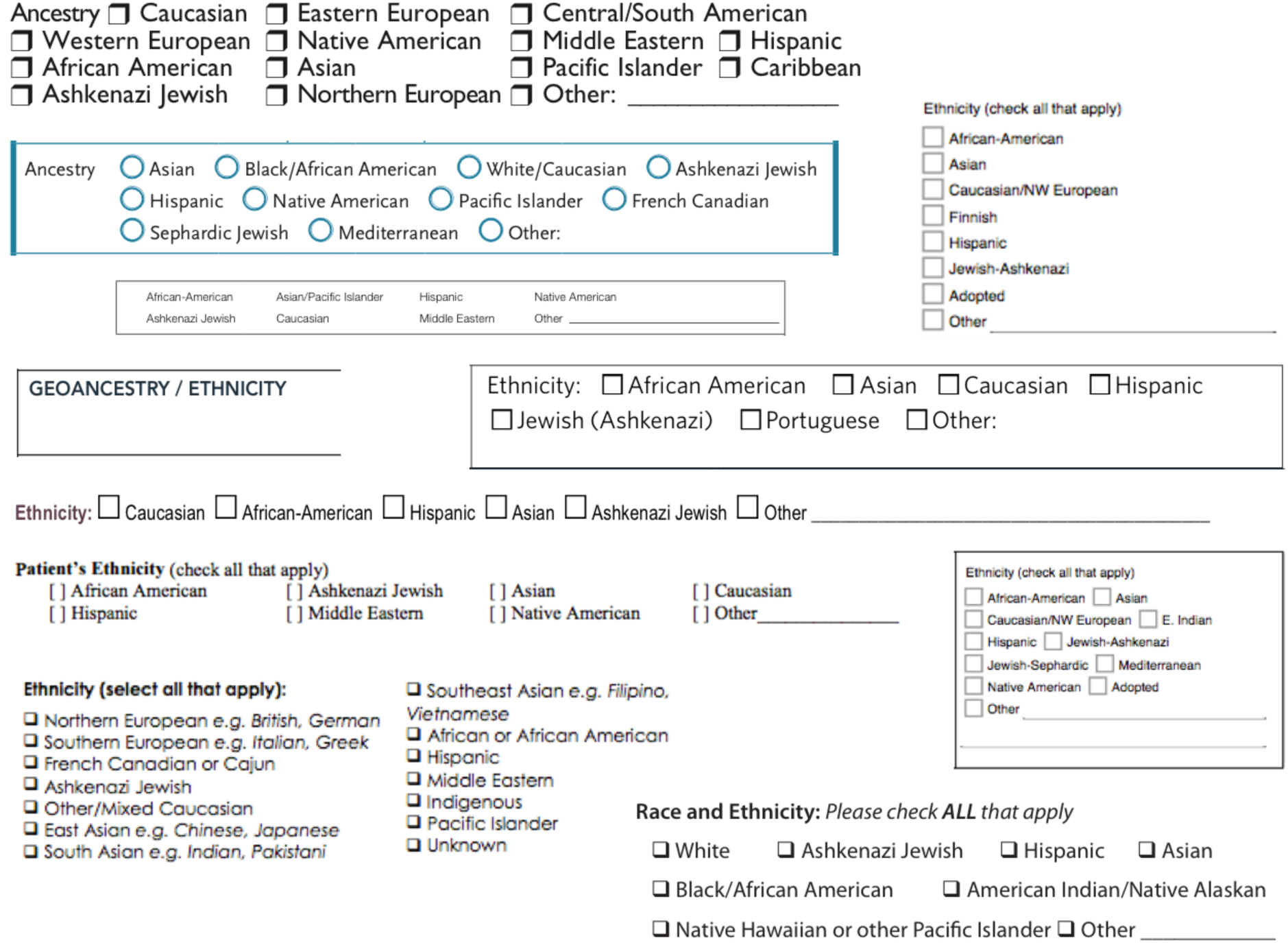
Screenshots of race, ethnicity, and ancestry questions on clinical laboratory requisition forms.

## Footnotes

1 https://www.clinicalgenome.org

2 https://clinvarminer.genetics.utah.edu

3 https://www.ncbi.nlm.nih.gov/medgen/

4 https://cser-consortium.org

5 https://allofus.nih.gov

6 https://grants.nih.gov/grants/guide/rfa-files/RFA-HG-16-011.html

7 https://www.genome.gov/10001691/nhgri-genome-sequencing-program-gsp/

8 https://www.nhlbi.nih.gov/science/trans-omics-precision-medicine-topmed-program

